# Genetic contributions to variation in human stature in prehistoric Europe

**DOI:** 10.1101/690545

**Authors:** Samantha L. Cox, Christopher B. Ruff, Robert M. Maier, Iain Mathieson

## Abstract

The relative contributions of genetics and environment to temporal and geographic variation in human height remain largely unknown. Ancient DNA has identified changes in genetic ancestry over time, but it is not clear whether those changes in ancestry are associated with changes in height. Here, we directly test whether changes over the past 38,000 years in European height predicted using DNA from 1071 ancient individuals are consistent with changes observed in 1159 skeletal remains from comparable populations. We show that the observed decrease in height between the Early Upper Paleolithic and the Mesolithic is qualitatively predicted by genetics. Similarly, both skeletal and genetic height remained constant between the Mesolithic and Neolithic and increased between the Neolithic and Bronze Age. Sitting height changes much less than standing height–consistent with genetic predictions–although genetics predicts a small Bronze Age increase that is not observed in skeletal remains. Geographic variation in stature is also qualitatively consistent with genetic predictions, particularly with respect to latitude. Finally, we hypothesize that an observed decrease in genetic heel bone mineral density in the Neolithic reflects adaptation to the decreased mobility indicated by decreased femoral bending strength. This study provides a model for interpreting phenotypic changes predicted from ancient DNA and demonstrates how they can be combined with phenotypic measurements to understand the relative contribution of genetic and developmentally plastic responses to environmental change.

**Significance:** Measurements of prehistoric human skeletal remains provide a record of changes in height and other anthropometric traits, over time. Often, these changes are interpreted in terms of plastic developmental response to shifts in diet, climate or other environmental factors. These changes can also be genetic in origin but, until recently, it has been impossible to separate the effects of genetics and environment. Here we use ancient DNA to directly estimate genetic changes in phenotypes and to identify changes driven not by genetics, but by environment. We show that changes over the past 35,000 years are largely predicted by genetics, but also identify specific shifts that are more likely to be environmentally driven.

## Introduction

Stature, or standing height, is one of the most heavily studied human phenotypes. It is easy to measure in living individuals and relatively straightforward to estimate from skeletal remains. As a consequence, geographic variation and temporal changes in stature are well documented (1–3), particularly in western Europe, where there is a comprehensive record of prehistoric changes (4). The earliest anatomically modern humans in Europe, present by 42-45,000 BP (5, 6), were relatively tall (mean adult male height in the Early Upper Paleolithic was ~174 cm). Mean male stature then declined from the Paleolithic to the Mesolithic (~164 cm) before increasing to ~167 cm by the Bronze Age (4, 7). Height can respond rapidly in a developmentally plastic manner to changes in environment, as demonstrated by large increases in Europe, and worldwide, during the secular trends of the 19^th^ and 20^th^ centuries (1, 4). In European countries today, mean adult male height is ~170-180 cm (1). It is broadly agreed that prehistoric changes were likely to have been driven by a combination of environmental (e.g. climate or diet) and genetic factors including drift, admixture and selection (4, 7–9), although the effects of these variables cannot be separated based on skeletal data alone. In this study, by combining the results of genome-wide association studies (GWAS) with ancient DNA, we directly estimate the genetic component of stature and test whether population-level skeletal changes between ~35,000 and 1,000 BP are consistent with those predicted by genetics.

Height is highly heritable (10–14), and therefore amenable to genetic analysis by genome-wide association studies (GWAS). With sample sizes of hundreds of thousands of individuals, GWAS have identified thousands of genomic variants that are significantly associated with the phenotype (15–17). Though the individual effect of each of these variants is tiny (on the order of +/− 1-2mm per variant (18)), their combination can be highly predictive. Polygenic risk scores (PRS) constructed by summing together the effects of all height-associated variants carried by an individual can now explain upwards of 30% of the phenotypic variance in populations of European ancestry (16). In effect, the PRS can be thought of as an estimate of “genetic height” that predicts phenotypic height, at least in populations closely related to those in which the GWAS was performed. One major caveat is that the predictive power of PRS is much lower in other populations (19). The extent to which differences in PRS between populations are predictive of population-level differences in phenotype is currently unclear (20). Recent studies have demonstrated that such differences may partly be artifacts of correlation between environmental and genetic structure in the original GWAS (21, 22). These studies also suggested best practices for PRS comparisons, including the use of GWAS summary statistics from large homogenous studies (instead of meta-analyses), and replication of results using summary statistics derived from within-family analyses that are robust to population stratification.

Bearing these caveats in mind, PRS can be applied to ancient populations thanks to recent technological developments that have dramatically increased ancient DNA (aDNA) sample sizes. These have provided remarkable insights into the demographic and evolutionary history of both modern and archaic humans across the world (23–25), particularly in Europe, and allow us to track the evolution of variants underlying phenotypes ranging from pigmentation to diet (26–29). In principle, PRS applied to ancient populations could similarly allow us to make inference about the evolution of complex traits. A few studies have used PRS to make predictions about the relative statures of ancient populations (29–31) but looked at only a few hundred samples in total and did not compare their predictions with stature measured from skeletons. Here, we compare measured skeletal data to genetic predictions and directly investigate the genetic contribution to height independent of environmental effects acting during development.

## Results

### PRS and skeletal measurements

We collected published aDNA data from 1071 ancient individuals from Western Eurasia (west of 50° E), dated to between 38,000 and 1100 years before present (BP) (27, 29, 30, 32–57). Using GWAS summary statistics for height from the UK Biobank (generated and made available by the Neale lab: http://nealelab.is/), we computed height PRS for each individual, using a P-value cutoff of 10^−6^, clumping variants in 250kb windows, and replacing missing genotypes with the mean across individuals (Methods). We refer to this as PRS(GWAS). Because of concerns about GWAS effect sizes being inflated by residual population stratification, we also computed a PRS where we used GWAS P-values to select SNPs, but computed the PRS using effect sizes estimated from a within-family test for ~17,000 sibling pairs from UK Biobank (Methods) which we refer to as PRS(GWAS/Sibs), and which should be unaffected by stratification. We also obtained stature estimates from 1159 individuals dating to between 33,700 and 1100 BP taken from a larger dataset of 2177 individuals with stature and body proportion estimates from substantially complete skeletons (4, 58). There is limited overlap in these datasets (12 individuals), but they cover the same time periods and broadly the same geographic locations (Supplementary Fig. 1), although the genetic data contain more individuals from further east (30-50° E) compared to the skeletal data. We divided these individuals into five groups based on date: Early Upper Paleolithic (>25,000 BP; EUP), Late Upper Paleolithic (25,000-11,000 BP; LUP), Mesolithic (11,000-5500 BP), Neolithic (8500-3900 BP) and post-Neolithic (5000-1100 BP, including the Copper and Bronze Ages, plus later periods), resolving individuals in the overlapping periods using either archaeological or genetic context (Methods). These groups broadly correspond to transitions in both archaeological culture and genetic ancestry (33, 38, 59) (Supplementary Fig. 1c-d, Supplementary Table 1).

### Trends in PRS for height are largely consistent with trends in skeletal stature

Both PRS and skeletal stature decreased from the EUP to Mesolithic periods and increased between the Neolithic and post-Neolithic (Supplementary Fig. 2). Fitting group (time period) as a covariate, we found a significant effect on PRS(GWAS) (ANOVA P= 1.9×10^−9^), PRS(GWAS/Sibs) (P=0.045) and skeletal stature (P=2.8×10^−11^). There was no evidence of difference between LUP, Mesolithic and Neolithic groups (Supplementary Fig. 3a-b), so we merged these three groups (we refer to the merged group as LUP-Neolithic). We find that PRS(GWAS) in the LUP-Neolithic period is 0.47 standard deviations (SD) lower than in the EUP (P=0.002), and 0.40 SD lower (P= 8.7×10^−11^) than in the post-Neolithic period (Fig. 1a). PRS(GWAS/Sib) shows a very similar pattern (Fig. 1b), demonstrating that this is not a result of differential relatedness of the ancient individuals to the structured present-day GWAS populations. Skeletal stature shows a qualitatively similar pattern to the genetic predictions (Fig. 1c), with a 1.5 SD (9.6cm; P=2.9×10^−7^) difference between EUP and LUP-Neolithic and a 0.27 SD (1.8cm; P=3.6×10^−5^) difference between LUP-Neolithic and post-Neolithic. Broad patterns of change in stature over time are therefore consistent with genetic predictions.

**Figure 1:**
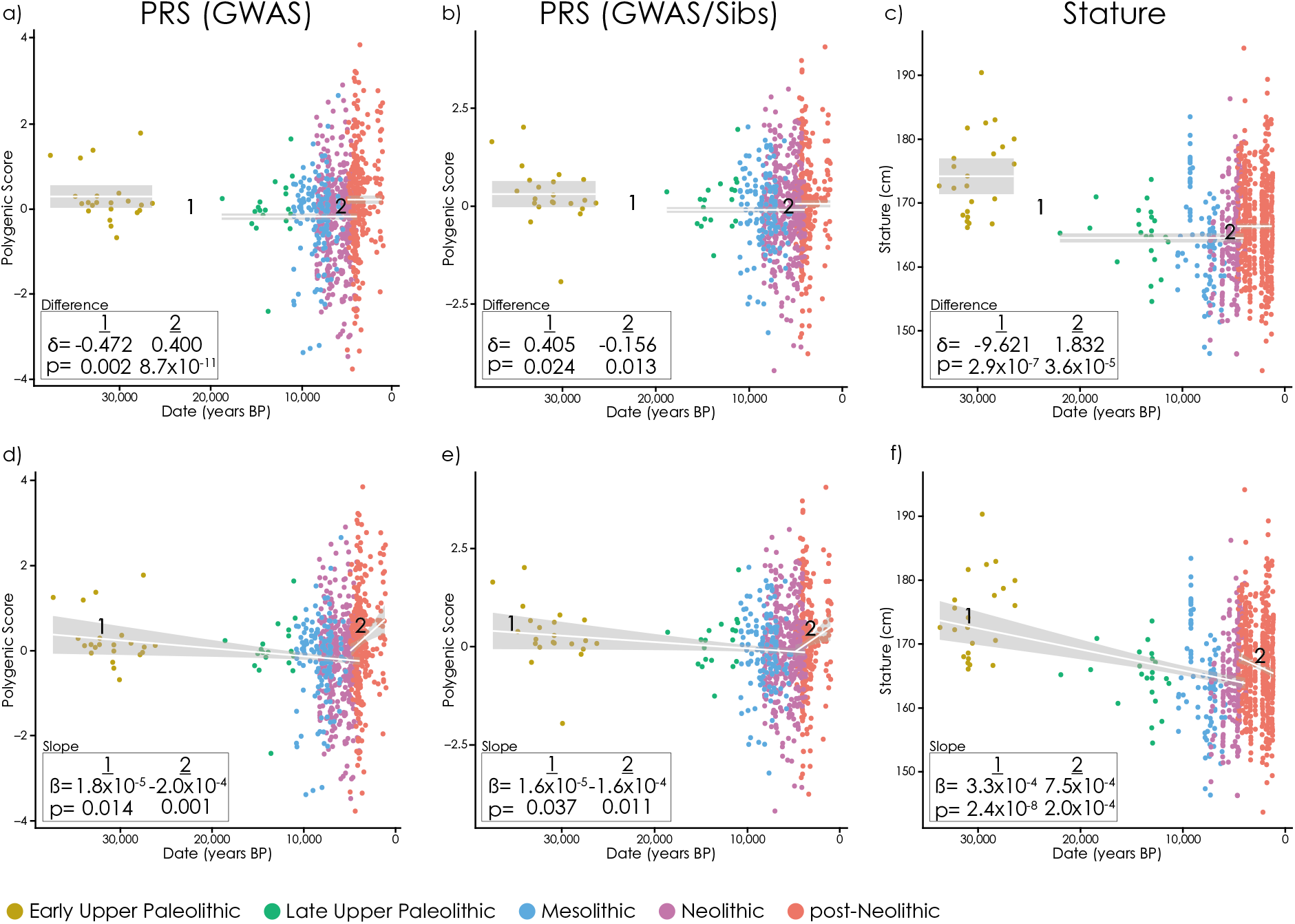
Changes in height PRS and stature though time. Each point is an ancient individual, white lines show fitted values, grey area is the 95% confidence interval, and boxes show parameter estimates and p-values for difference in means (δ) and slopes (β). **a-c**) PRS(GWAS) (a), PRS(GWAS/Sibs) (b) and skeletal stature (c) with constant values in the EUP, LUP-Neolithic and post-Neolithic. **d-e**) PRS(GWAS) (d), PRS(GWAS/Sibs) (e) and skeletal stature (f) showing a linear trend between EUP and Neolithic and a different trend in the post-Neolithic.

Additionally, we fit a piecewise linear model allowing PRS to decrease from the EUP to the Neolithic and then increase and change slope in the post-Neolithic (Fig. 1d-f). In this model, PRS(GWAS) decreases by about 1.8×10^−5^ SD/year (P=0.014) from EUP to Neolithic, and increases by 2.0×10^−4^ SD/year (P=0.001) post-Neolithic (Fig. 1d). PRS(GWAS/sib) decreases by about 1.6×10^−5^ SD/year (P=0.037) from EUP to Neolithic, then increases by 1.6×10^−4^ SD/year throughout the period (P=0.011; Fig. 1e). Again, these changes are qualitatively consistent with changes in stature (Fig. 1f), with a 4.7×10^−5^ SD/year (3.3×10^−4^ cm/year; P=2.4×10^−8^) decrease from EUP to Mesolithic, and an increase of ~0.5 SD into the Neolithic. However, in this model stature, unlike PRS, actually decreases during the post-Neolithic period (7.5×10^−4^ cm/year; P=2.0×10^−4^).

To further explore these trends, we fitted a broader range of piecewise linear models to both datasets (Methods; Supplementary Table 1; Supplementary Fig. 4-6). In the most general model we allowed both the mean and the slope of PRS or stature with respect to time to vary between groups. More constrained models fix some of these parameters to zero–eliminating change over time–or merging two adjacent groups. We compared the fit of these nested models using Akaike’s Information Criterion (AIC, Supplementary Table 2). The linear model in Fig. 1d-f is one of the best models in this analysis. In general, all the best-fitting models support the pattern–for both PRS and measured stature–of a decrease between the EUP and Mesolithic and an increase between the Neolithic and post-Neolithic (Supplementary Fig. 4-6). Some models suggest that the increase in stature–but not PRS–may have started during the Neolithic (Supplementary Fig. 6a-c). Finally, we confirmed that these results were robust to different constructions of the PRS–using 100kb and 500kb clustering windows rather than 250kb (Supplementary Fig. 7-8).

### Sitting height PRS is partially consistent with trends in body proportions

Standing height is made up of two components: leg length and sitting height (made up of the length of the trunk, neck and head), with a partially overlapping genetic basis (60). Throughout European prehistory, changes in leg length tended to be larger than changes in sitting height (4). We constructed PRS(GWAS) and PRS(GWAS/Sibs) for sitting height and analyzed them in the same way as standing height (Fig. 2). In contrast to standing height, we find no evidence of change between the EUP and Neolithic. Both PRS(GWAS) and PRS(GWAS/Sibs) do increase, either between the Neolithic and post-Neolithic, or during the post-Neolithic period (Fig. 2a,b,d & e). On the other hand, using only skeletons with complete torsos to estimate sitting height, we find no evidence of change in any period. Thus, the skeletal data are consistent with the genetic data for the EUP-Neolithic period, but inconsistent in the post-Neolithic period, where PRS predicts an increase that is not reflected in the skeletons. This could be because of more limited skeletal measurements (only 236 out of 1159 skeletons are sufficiently complete to estimate sitting height directly), because the change in PRS is artefactual, it is being buffered by non-genetic effects, or by opposing genetic effects which we do not capture. Overall, we find mixed consistency between PRS and skeletal measurements. The decrease in standing but not sitting height between the EUP and Neolithic is consistent in both, as is the increase in standing height between the Neolithic and post-Neolithic. However, PRS predicts a continued increase in stature through the post-Neolithic period that is not seen in skeletal remains.

**Figure 2:**
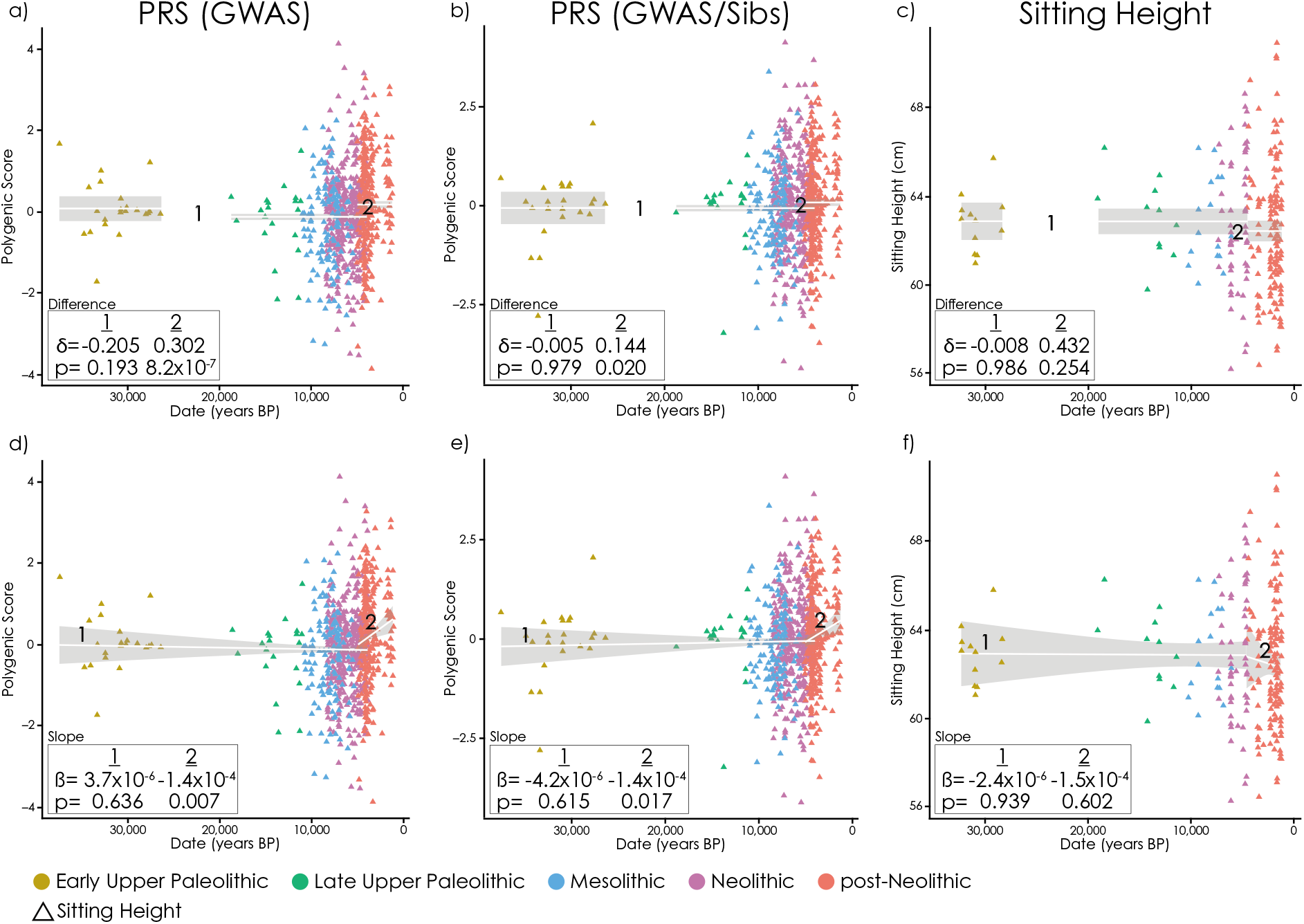
Changes in sitting height PRS and sitting height though time. Each point is an ancient individual, lines show fitted values, grey area is the 95% confidence interval, and boxes show parameter estimates and p-values for difference in means (δ) and slopes (ß). **a-c**) PRS(GWAS) (a), PRS(GWAS/Sibs) (b) and skeletal sitting height, with constant values in the EUP, LUP-Neolithic and post-Neolithic. **d-e**) PRS(GWAS) (d), PRS(GWAS/Sibs) (e) and skeletal sitting height showing a linear trend between EUP and Neolithic and a different trend in the post-Neolithic.

### Geographic variation in standing height

As well as varying through time, human stature is stratified by geography, with trends related to both longitude and latitude (61). North-South trends following Allen’s (62) and Bergmann’s (63) rules are most often interpreted as environmental adaptations to the polar-equatorial climate gradient. Today, Northern Europeans are generally taller than Southern Europeans (1), a pattern which emerged between the Mesolithic and post-Neolithic (4, 7). Longitudinal variation within Europe is present during the Mesolithic (64), though these trends are difficult to interpret due to sampling bias across the time period (4). We therefore tested whether geographic variation in PRS could explain these geographic trends, as it partially explains temporal trends.

We regressed the residuals from our fitted linear height model (the model shown in Fig. 1d-f) on longitude and latitude. Stature increases significantly with latitude (P=1.2×10^−10^) in the post-Neolithic period. PRS(GWAS) increases in the post-Neolithic (P=0.006) although this is not replicated by PRS(GWAS/Sibs) (P=0.557). PRS does not increase significantly with latitude in the EUP-Neolithic period. There is some evidence of a modest trend in stature in the EUP-Neolithic period (Fig. 3c). However, there is only evidence for this in the Neolithic, not in the EUP-Mesolithic (Supplementary Fig. 9a). Further, because time and geography are correlated in our Neolithic sample, this can also be explained by a temporal increase during the Neolithic, in which case there is no geographic trend (Supplementary Fig. 9b).

**Figure 3:**
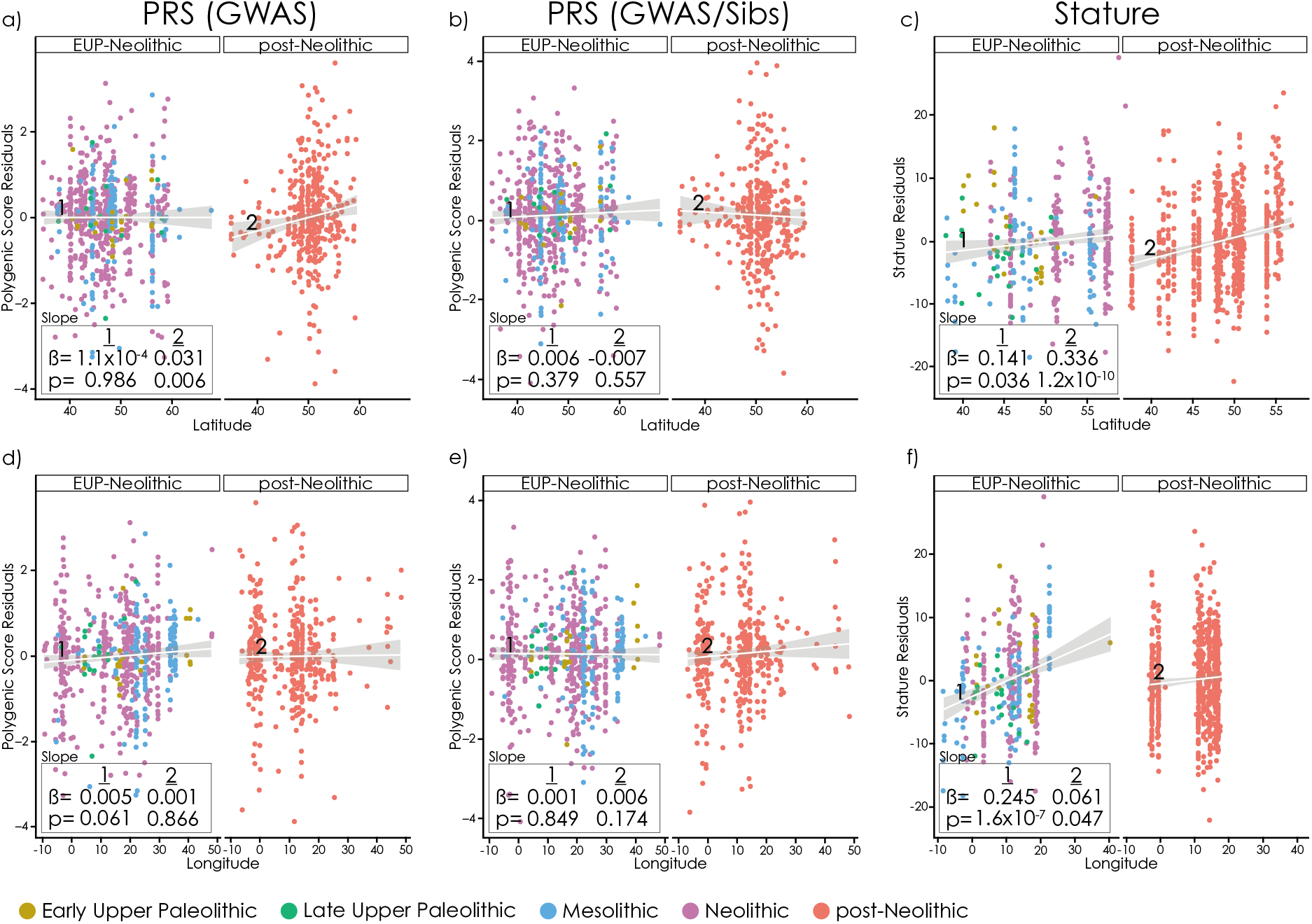
Geographic variation in PRS and skeletal standing height. Residuals for the linear height model (Fig. 1 d-f) against **a-c**) latitude and **d-f**) longitude. Each point is an ancient individual, lines show fitted values, grey area is the 95% confidence interval, and boxes show parameter estimates (β) and p-values for slopes.

In contrast to latitude, there is a significant increasing trend of stature with longitude before but not during the Neolithic (0.36 cm/degree P=1.6×10^−7^; Fig. 3, Supplementary Fig. 9c). This is partly driven by a small number of samples from a single site, but still persists if these samples are removed (0.20 standardized residuals per degree, P=0.004; Supplementary Fig. 9d). There is little or no trend (0.06 cm/degree; P=0.047) in the post-Neolithic period (Fig. 3f). We find no evidence for longitudinal clines in PRS. In summary, we find that stature increases with latitude in the post-Neolithic, possibly in the Neolithic, but not before. This cline may have a genetic basis. Stature also increases with longitude, particularly in the Mesolithic, but this cline is not predicted by genetics.

### Correlated changes in bone density PRS and femoral bending strength

Beyond stature, we wanted to investigate the utility of using PRS to interpret other measurable phenotypes in ancient individuals. Decreased mobility though time, associated with large-scale lifestyle transitions between hunting-gathering, agriculture, and ultimately modern industrialism, is well documented through declines in lower limb bone diaphyseal strength and trabecular density (4, 65, 66). Today, heel bone mineral density (hBMD) is often used as an indicator of general activity levels in younger people (67) and of osteoporosis in older individuals (68, 69); UK Biobank has GWAS data for this trait, indirectly estimated by ultrasound. However, evaluating differences in BMD in archaeological and paleontological specimens can be problematic. In the short term soil leaches bone minerals, while later the bone begins to fossilize, leading to unpredictable patterns of density in ancient remains (70) and requiring special processing methods (65) that are difficult to apply to large samples. However, femoral diaphyseal bending strength can be calculated from bone cross-sectional geometric measurements that are not as affected by bone preservation (71). Here we focus on anteroposterior bending strength (section modulus) of the midshaft femur (FZx), which has been linked specifically to mobility (72). Since both trabecular density and diaphyseal strength should respond to mobility and activity levels, we reasoned that they would be likely to show correlated patterns of temporal change. Following established protocols (71), we standardized FZx first by sex, then the product of estimated body mass and femoral length (4). Qualitatively, PRS(GWAS) and FZx show similar patterns, decreasing through time (Fig. 4, Supplementary Fig. 2-3). There is a significant drop in FZx (Figure 4c) from the Mesolithic to Neolithic (P= 1.2×10^−8^) and again from the Neolithic to post-Neolithic (P=1.5×10^−13^). PRS(GWAS) for hBMD decreases significantly from the Mesolithic to Neolithic (Fig. 4a; P=5.5×10^−12^), which is replicated in PRS(GWAS/Sibs) (P=7.2×10^−10^; Fig. 4b); neither PRS shows evidence of decrease between the Neolithic and post-Neolithic. We hypothesize that both FZx and hBMD responded to the reduction in mobility that accompanied the adoption of agriculture (72). In particular, the lower genetic hBMD and skeletal FZx of Neolithic compared to Mesolithic populations may represent adaptation to the same change in environment although we do not know the extent to which the change in FZx was driven by genetic or plastic developmental response to environmental change. On the other hand, FZx continues to decrease between the Neolithic and post-Neolithic (Fig. 4c,f)–which is not reflected in the hBMD PRS (Fig. 4 a-b,d-e). One possibility is that the two phenotypes responded differently to the post-Neolithic intensification of agriculture. Another is that the non-genetic component of hBMD, which we do not capture here, also continued to decrease.

**Figure 4:**
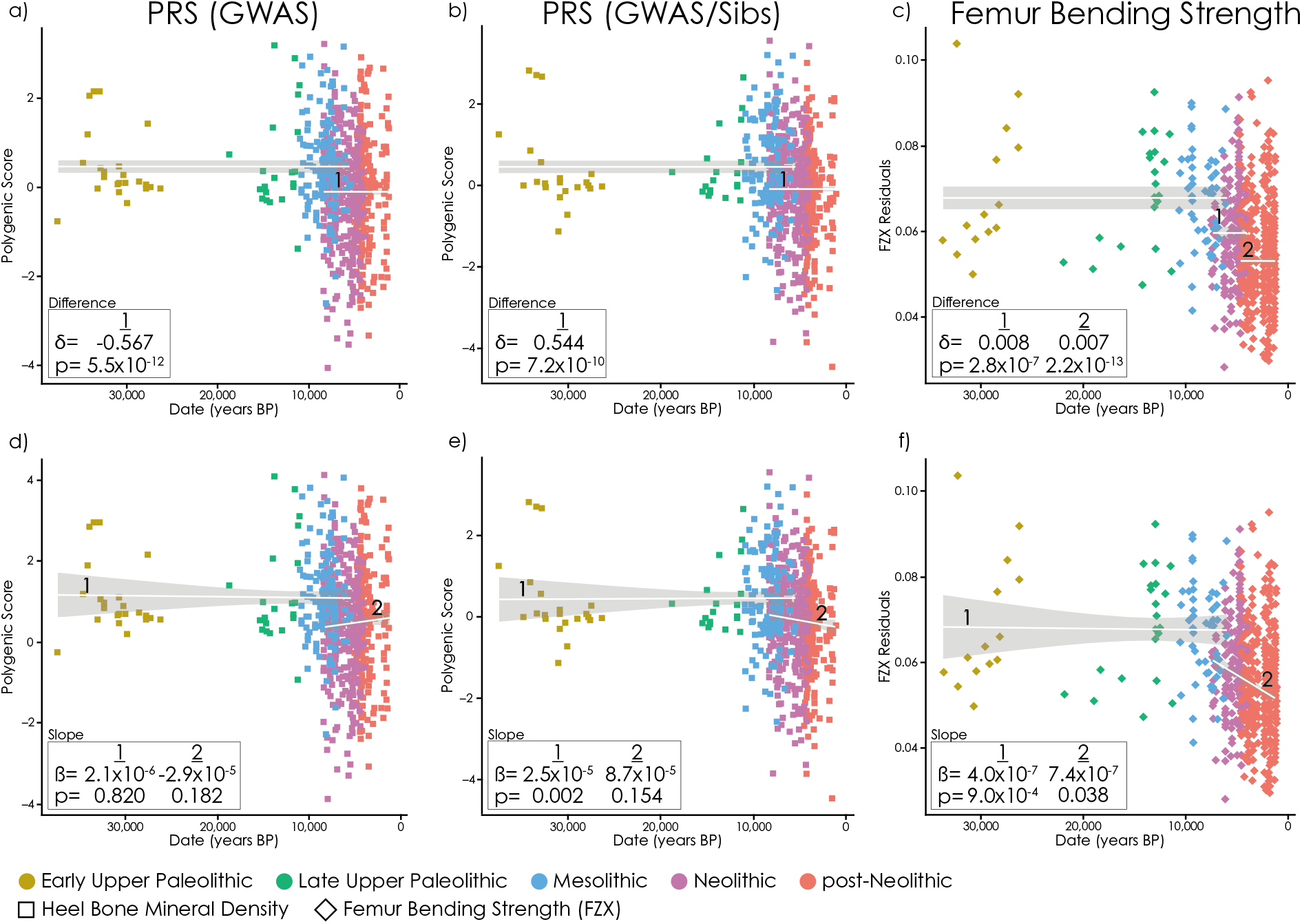
Changes in heel bone mineral density (hBMD) PRS and femur bending strength (FZx) though time. Each point is an ancient individual, lines show fitted values, grey area is the 95% confidence interval, and boxes show parameter estimates and p-values for difference in means (δ) and slopes (β). **a-b**) PRS(GWAS) (a) and PRS(GWAS/Sibs) (b) for hBMD, with constant values in the EUP-Mesolithic and Neolithic-post-Neolithic. **c**) FZx constant in the EUP-Mesolithic, Neolithic and post-Neolithic. **d-e**) PRS(GWAS) (d) and PRS(GWAS/Sibs) (e) for hBMD showing a linear trend between EUP and Mesolithic and a different trend in the Neolithic-post-Neolithic. **f**) FZx with a linear trend between EUP and Mesolithic and a different trend in the Neolithic-post-Neolithic.

### Are changes in PRS driven by selection or genetic drift?

The *Q_x_* statistic (73) can be used to test for polygenic selection. We computed it for increasing numbers of SNPs from each PRS (Fig. 5a-c), between each pair of adjacent time periods and over all time periods. We estimated empirical P-values by replacing allele frequencies with random derived allele frequency-matched SNPs from across the genome, while keeping the same effect sizes. To check these *Q_x_* results, we simulated a GWAS from the UK Biobank dataset (Methods), then used these effect sizes to compute simulated *Q_x_* statistics. The *Q_x_* test suggests selection between the Neolithic and Post-Neolithic for stature (P<1 ×10^−4^; Fig. 5a), which replicates using effect sizes estimated within siblings (10^−4^<P<10^−2^; Supplementary Fig. 10). The reduction in the sibling effect compared to the GWAS effect sizes is consistent with the reduction expected from the lower sample size (Supplementary Figure 10). However, several (11/200) simulated datasets produce higher *Q_x_* values than observed in the real data (Fig. 5d). This suggests that re-estimating effect sizes between siblings may not fully control for the effect of population structure and ascertainment bias on the *Q_x_* test. The question of whether selection contributes to the observed differences in height PRS remains unresolved.

**Figure 5:**
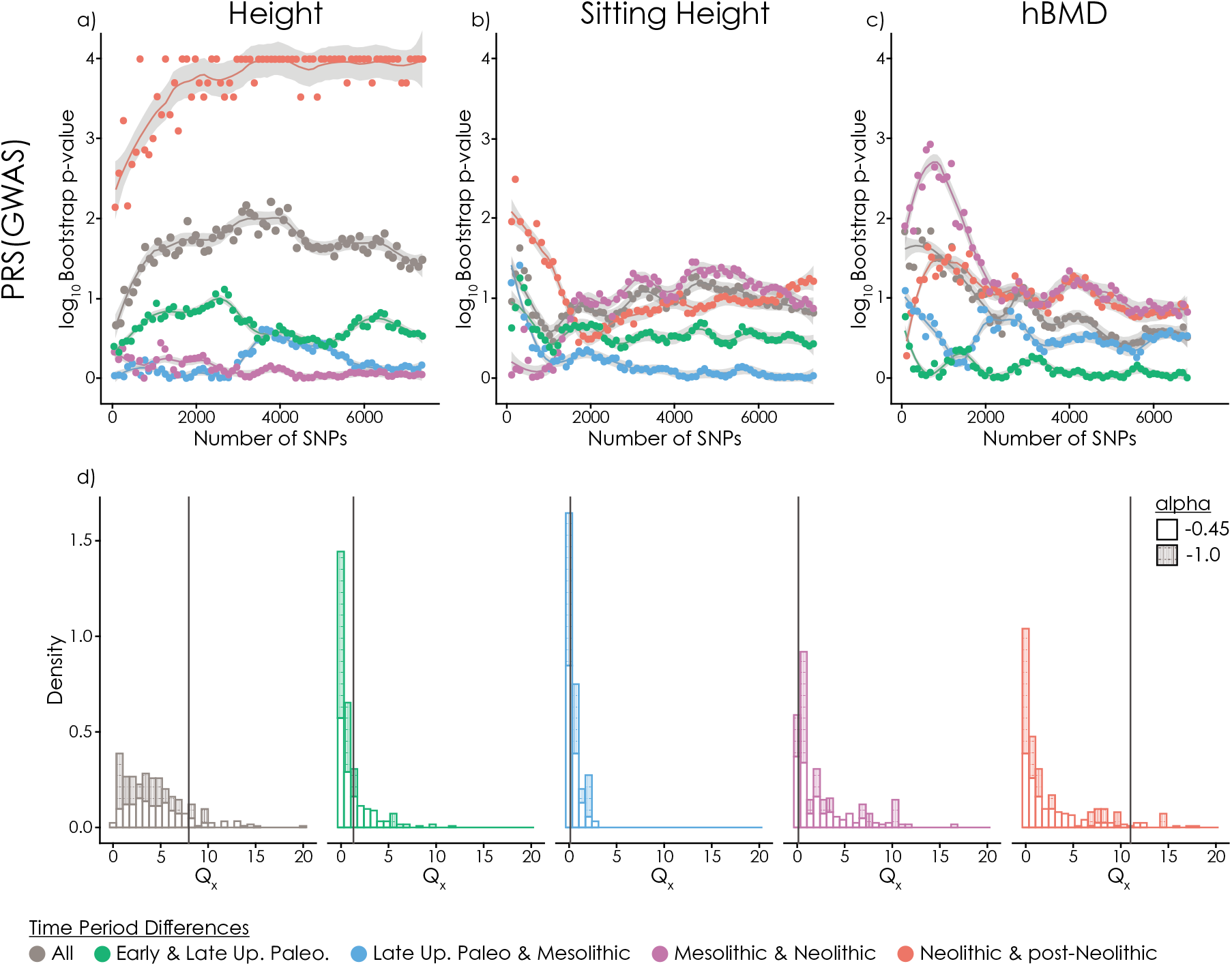
Signals of selection on standing height, sitting height and bone mineral density. **a-c**) Log_10_ bootstrap P-values for the *Q_x_* statistics (y-axis, capped at 4) for GWAS signals. We tested each pair of adjacent populations, and the combination of all of them (“All”). We ordered PRS SNPs by increasing P-value and tested the significance of *Q_x_* for increasing numbers of SNPs (x-axis). **d**) Distribution of *Q_x_* statistics in simulated data (Methods). Observed height values for 6800 SNPs shown by vertical lines.

For sitting height, we find little evidence of selection in any time period (P<10^−2^) We conclude that there was most likely selection for increased standing but not sitting height in the Steppe ancestors of Bronze Age European populations, as previously proposed (29). One potential caveat is that, although we re-estimated effect sizes within siblings, we still used the GWAS results to identify SNPs to include. This may introduce some subtle confounding, which remains a question for future investigation. Finally, using GWAS effect sizes, we identify some evidence of selection on heel BMD when comparing Mesolithic and Neolithic populations (10^−3^<P<10^−2^; Fig. 5c). However, this signal is relatively weak when using within-sibling effect sizes, and disappears when we include more than about 2000 SNPs.

## Discussion

We showed that the well-documented temporal and geographic trends in stature in Europe between the Early Upper Paleolithic and the post-Neolithic period are broadly consistent with those that would be predicted by polygenic risk scores (PRS) computed using present-day GWAS results combined with ancient DNA. However, because of the limited predictive power of current PRS, we cannot provide a quantitative estimate of how much of the variation in phenotype between populations might be explained by variation in PRS. Similarly, we cannot say whether the changes were continuous, reflecting evolution through time, or discrete, reflecting changes associated with known episodes of replacement or admixture of populations that have diverged genetically over time. Finally, we find cases where predicted genetic changes are discordant with observed phenotypic changes–emphasizing the role of developmental plasticity in response to environmental change and the difficulty in interpreting differences in PRS in the absence of phenotypic data.

Our results indicate two major episodes of change in genetic height. First, there was a reduction in standing height PRS–but not sitting height PRS–between the Early and Late Upper Paleolithic, coinciding with an almost total population replacement (33). These genetic changes are consistent with the decrease in stature–driven by leg length–observed in skeletons during this time period (4, 64, 74, 75). One possibility is that the stature decrease in the ancestors of the LUP populations could have been adaptive, driven by changes in resource availability (76) or to a colder climate (61). Comparison between patterns of phenotypic and genetic variation suggest that, on a broad scale, variation in body proportions among present-day people reflects adaptation to environment largely along latitudinal gradients (77, 78). Early Upper Paleolithic populations in Europe would have migrated relatively recently from more southern latitudes and had body proportions that are typical of present-day tropical populations (75). The populations that replaced them would have had more time to adapt to the colder climate of northern latitudes. On the other hand, we do not find genetic evidence for selection on stature during this time period–suggesting that the changes may have been neutral and not adaptive.

The second episode of change in height is either between the Neolithic and post-Neolithic, or during the post-Neolithic period. This period is characterized by the eastward movement of substantial amounts of “Steppe ancestry” into Central and Western Europe (27, 30, 38, 50). Our results are thus consistent with previous results that migration and admixture from Bronze Age populations of the Eurasian steppe increased genetic height in Europe (29, 30). Whether this increase was driven by selection in the ancestors of these populations remains unresolved. The geographic gradient of increasing skeletal stature is unclear in the Paleolithic, largely West-East in the Mesolithic (7, 64) and largely South-North by the Bronze Age (4, 7, 9). Latitudinal, but not longitudinal, patterns are qualitatively consistent with geographic patterns in PRS suggesting that, like temporal variation, both genetics and environment contribute to geographic variation.

A major confounding factor in analysis of temporal and geographic variation in PRS, particularly in the Bronze Age, is genetic population structure. Present-day European population structure is correlated with geography and largely driven by variation in Steppe ancestry proportion, with more Steppe ancestry in Northern Europe and less in Southern Europe (38). Suppose that environmental variation in stature is also correlated with geography, and that Northern Europeans are taller than Southern Europeans for entirely non-genetic reasons. Then, GWAS that do not completely correct for stratification will find that genetic variants that are more common in Steppe populations than Neolithic populations are associated with increased height. When these GWAS results are used to compute PRS for ancient populations, they will predict that Steppe ancestry populations were genetically taller simply because they are more closely related to present-day Northern Europeans (21, 22). In this study, we attempted to avoid this confounding in two ways: first, by using GWAS effect sizes from the UK Biobank–a homogenous dataset that should be well-controlled for population stratification, and second, by replicating our results after re-estimating the effect sizes within siblings, which should be robust to population stratification, though less precise. However, we cannot exclude the possibility that some confounding remains, for example because although we re-estimated effect sizes using the within-siblings design, we still ascertained loci using the GWAS results. A related concern is that, even in the absence of bias, variance explained by the PRS is likely to decrease as we move back in time and ancient populations become less closely related to present-day populations (19). However, our results indicate that the PRS still captures enough of the genetic variance to predict changes in phenotype even though, presumably, we would gain even more resolution using a PRS that captured more of the genetic variation in the phenotype.

As well as genetic contributions to phenotype, our results shed light on possible environmental contributions. In some cases, we can make hypotheses about the relationship between environmental or lifestyle changes, and genetic change. For example, if we interpret change in femur bending strength as reflecting a decrease in mobility, the coincident Mesolithic/Neolithic change in heel bone mineral density PRS can be seen as a genetic response to this change. However, in the Neolithic/post-Neolithic periods, the two observations are decoupled. This emphasizes the role of developmental plasticity in response to changes in environment, and of joint interpretation of phenotypic and genetic variables. Even when looking at the same phenotype, we find cases where genetic predictions and phenotypic data are discordant–for example in post-Neolithic sitting height. We must therefore be cautious in the interpretation of predicted genetic patterns where phenotypes cannot be directly measured, even if it is possible to control stratification. Predicted genetic changes should be used as a baseline, against which non-genetic effects can be measured and tested.

## Methods

### Ancient DNA and polygenic risk score construction

We collected published ancient DNA data from 1122 ancient individuals, taken from 29 publications. The majority of these individuals had been genotyped using an in-solution capture reagent (“1240k”) that targets 1.24 million single nucleotide polymorphisms (SNPs) across the genome. Because of the low coverage of most of these samples, the genotype data are pseudo-haploid. That is, there is only a single allele present for each individual at each site, but alleles at adjacent sites may come from either of the two chromosomes of the individual. For individuals with shotgun sequence data, we selected a single read at each 1240k site. We obtained the date of each individual from the original publication. Most of the samples have been directly radiocarbon dated, or else are securely dated by context. We summarized the genetic relationships between ancient and present-day groups by computing *F_ST_* using smartpca v16000 (79) (Supplementary Table 1), multidimensional scaling using pairwise distances computed using plink v1.90b5.3 (options --distance flat-missing 1-ibs) (80) (Supplementary Fig. 1c) and unsupervised ADMXITURE (81) (Supplementary Fig. 1d).

We obtained GWAS results from the Neale lab UK Biobank page (http://www.nealelab.is/uk-biobank/; Round 1, accessed February and April 2018). To compute PRS, we first took the intersection of the 1240k sites and the association summary statistics. We then selected a list of SNPs to use in the PRS by selecting the SNP with the lowest P-value, removing all SNPs within 250kb, and repeating until there were no SNPs remaining with P-value less than 10^−6^. We then computed PRS for each individual by taking the sum of genotype multiplied by effect size for all included SNPs. Where an individual was missing data at a particular SNP, we replaced the SNP with the average frequency of the SNP across the whole dataset. This has the effect of shrinking the PRS towards the mean and should be conservative for the identification of differences in PRS. We confirmed that there was no correlation between missingness and PRS, to make sure that missing data did not bias the results (correlation between missingness and PRS ρ=0.02; P=0.44, Supplementary Fig. 11). Finally, we normalized the PRS across individuals to have mean 0 and standard deviation 1.

We estimated within-family effect sizes from 17,358 sibling pairs in the UK Biobank to obtain effect estimates that are unaffected by stratification. Pairs of individuals were identified as siblings if estimates of IBS0 were greater than 0.0018 and kinship coefficients were greater than 0.185. Of those pairs, we only retained those where both siblings were classified by UK Biobank as “white British”, and randomly picked two individuals from families with more than two siblings. We used Hail (82) to estimate within-sibling pair effect sizes for 1,284,881 SNPs by regressing pairwise phenotypic differences between siblings against the difference in genotype. We included pairwise differences of sex (coded as 0/1) and age as covariates, and inverse-rank-normalized the phenotype before taking the differences between siblings. To combine the GWAS and sibling results, we first restricted the GWAS results to sites where we had estimated a sibling effect size and replaced the GWAS effect sizes by the sibling effects. We then restricted to 1240k sites and constructed PRS in the same way as for the GWAS results.

To test whether the differences in the GWAS and GWAS/Sibs PRS results can be explained by differences in power, we created subsampled GWAS estimates which matched the sibling in the expected standard errors, by determining the equivalent sample size necessary and randomly sampling *N_sub_* individuals. 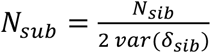 where *δ_sib_* is the difference in normalized phenotype between siblings after accounting for the covariates age and sex.

### Stature data

We obtained stature data from Ruff (2018) (4) (data file and notes available at http://www.hopkinsmedicine.org/fae/CBR.html), which also includes estimated body mass, femoral midshaft anteroposterior strength (FZx), and other osteometric dimensions. Statures and body masses were calculated from linear skeletal measurements using anatomical reconstruction or sample-specific regression formulae (4, 58). We calculated sitting height as basion-bregma (cranial) height (BBH) plus vertebral column length (VCL). Analysis was restriced to 1159 individuals dated earlier than 1165 BP (651 males and 508 females), of which 1130 had estimates for stature, 1014 for FZx and 236 for sitting height. Sitting and standing height were standardized for sex by adding the mean difference between male and female estimates to all the female values. Sex differences in stature remain relatively constant over time (4), making it reasonable to adjust all female heights by the same mean value. Male/Female counts in each period were: EUP 13/10, LUP 15/8, Mesolithic 56/37, Neolithic 130/91 and post-Neolithic 437/362. For FZx we first standardized for sex as we did for stature then divided each by estimated body mass multiplied by biomechanical femur length (4).

### Grouping

We grouped individuals into broad categories based on date and, in some cases, archeological and genetic context. All individuals were assigned to one time period group, based on median age estimates of the sample obtained from the original publications. Date ranges for each time period are based on a combination of historical, climatic, and archaeological factors. The Early Upper Paleolithic comprises all samples older than 25,000 BP, which roughly coincides with the end of the last glacial maximum (LGM). The Late Upper Paleolithic begins when the European glaciers are beginning to recede (25,000 BP) and extends until 11,000 BP and a shift in lithic technology that is traditionally used to delineate the beginning of the Mesolithic period. Transitions between the Mesolithic, Neolithic, and Bronze Age are staggered throughout Europe, so creating universally applicable date ranges is not possible. We instead defined overlapping transition periods between the Mesolithic and Neolithic periods (8500-5500 BP) and between the Neolithic and post-Neolithic (5000-3900 BP). For the genetic data, samples in the overlapping periods were assigned based on genetic population affiliation, inferred using supervised ADMIXTURE (81) which, in most of Western Europe, corresponds closely to archaeological context (38, 48). In particular, the Mesolithic/Neolithic overlap was resolved based on whether each individual had more (Neolithic) or less (Mesolithic) than 50% ancestry related to northwest Anatolian Neolithic Farmers. The Neolithic/post-Neolithic overlap was resolved based on whether individuals had more than 25% ancestry related to Bronze Age Steppe populations (“Steppe ancestry”; See (83) for details). For the skeletal data, group assignment in the overlapping periods was determined by the archaeology of each site. Broadly, sites belonging to the Neolithic have transitioned to agricultural subsistence. Similarly, post-Neolithic populations are broadly defined by evidence of metal working (Copper, Bronze and Iron Ages, and later periods). In particular, we included Late Eneolithic (Copper Age) sites associated with Corded Ware and Bell Beaker material culture in the post-Neolithic category but for consistency with the genetic classifications, we included 8 Early Eneolithic (before 4500 BP) individuals in the Neolithic category, since this precedes the appearance of Steppe ancestry in Western Europe. We excluded samples more recent than 1165 BP.

### Linear models

We fitted a series of linear models to changes in both PRS and stature data with time. In the most general model, we allow both the intercept and slope to vary between groups. We then either force some of the slopes to be zero, or some of the adjacent groups to have identical parameters. We describe the models using underscores to indicate changes in parameters, lowercase to indicate slopes (change with respect to time) fixed to zero, and upper case to indicate free slopes (i.e. linear trends with time). For example, “E_L_M_N_B” is the most general model, “elmnb” indicates that all groups have the same mean and there is no change with time, and “ELMN_B” indicates that the first four groups share the same parameters, and the post-Neolithic has different parameters. The models shown in Figures 1 and 2 are “e_lmn_b” (panels a-b), “e_lm_nb” (panel c), “ELMN_B” (panels d-e) and “ELM_NB” (panel f). To analyze geographic variation, we used the residuals of the “ELMN_B” model for the PRS and “ELM_NB” for skeletal stature, and fitted regressions against latitude and longitude.

### Polygenic selection test

We computed bootstrap P-values for the *Q_x_* statistic (73) by re-computing the statistic for random sets of SNPs in matched 5% derived allele frequency bins (polarized using the chimpanzee reference gnome panTro2). For each bootstrap replicate, we keep the original effect sizes but replace the frequencies of each SNP with one randomly sampled from the same bin. Unlike the PRS calculations, we ignored missing data, since the *Q_x_* statistic uses only the population-level estimated allele frequencies and not individual-level data. We tested a series of nested sets of SNPs (x-axis in Fig. 5), adding SNPs in 100 SNP batches, ordered by increasing P-value, down to a P-value of 0.1.

### Simulated GWAS data

We simulated GWAS, generating causal effects at a subset of around 159,385 SNPs in the intersection of SNPs which passed QC in the UK Biobank GWAS, are part of the 1240k capture, and are in the POBI dataset (84). We assumed that the variance of the effect size of an allele of frequency *f* was proportional to *[f(1-f)]^α^*, where the parameter *a* measures the relationship between frequency and effect size (85). We performed 100 simulations with *α*=−1 (the most commonly used model where each SNP explains the same proportion of phenotypic variance) and 100 with *α*=−0.45 as estimated for height (85). We then added an equal amount of random noise to the simulated genetic values, so that the SNP heritability equaled 0.5. We tested for association between these SNPs and the simulated phenotypes. Using these results as summary statistics, we computed PRS and *Q_x_* tests using the pipeline described above.

## Acknowledgments

I.M. was supported by a Research Fellowship from the Alfred P. Sloan foundation, and a New Investigator Research Grant from the Charles E. Kaufman Foundation [KA2018-98559]. Skeletal data were collected in collaboration with Brigitte Holt, Markku Niskanen, Vladimir Sladék, and Margit Bernor, with the support of the National Science Foundation [BCS-0642297 and BSC-0642710] and the Grant Agency of the Czech Republic and the Academy of Finland and Finnish Cultural Foundation. We thank Jeremy Berg and Eva Rosenstock for helpful comments on an earlier version of the manuscript. This research made use of the UK Biobank Resource under Application Number 33923. The project was initially conceived during discussions at the workshop “Human stature in the Near East and Europe in a long-term perspective” at the Freie Universität Berlin 25-27 April 2018, organized as part of the Emmy-Noether-Projekt “LiVES” funded by the German Research Foundation, Grant Nr. RO4148/1 (PI Eva Rosenstock).

