## Supplementary Figures for "Genetic contributions to variation in human stature in prehistoric Europe"

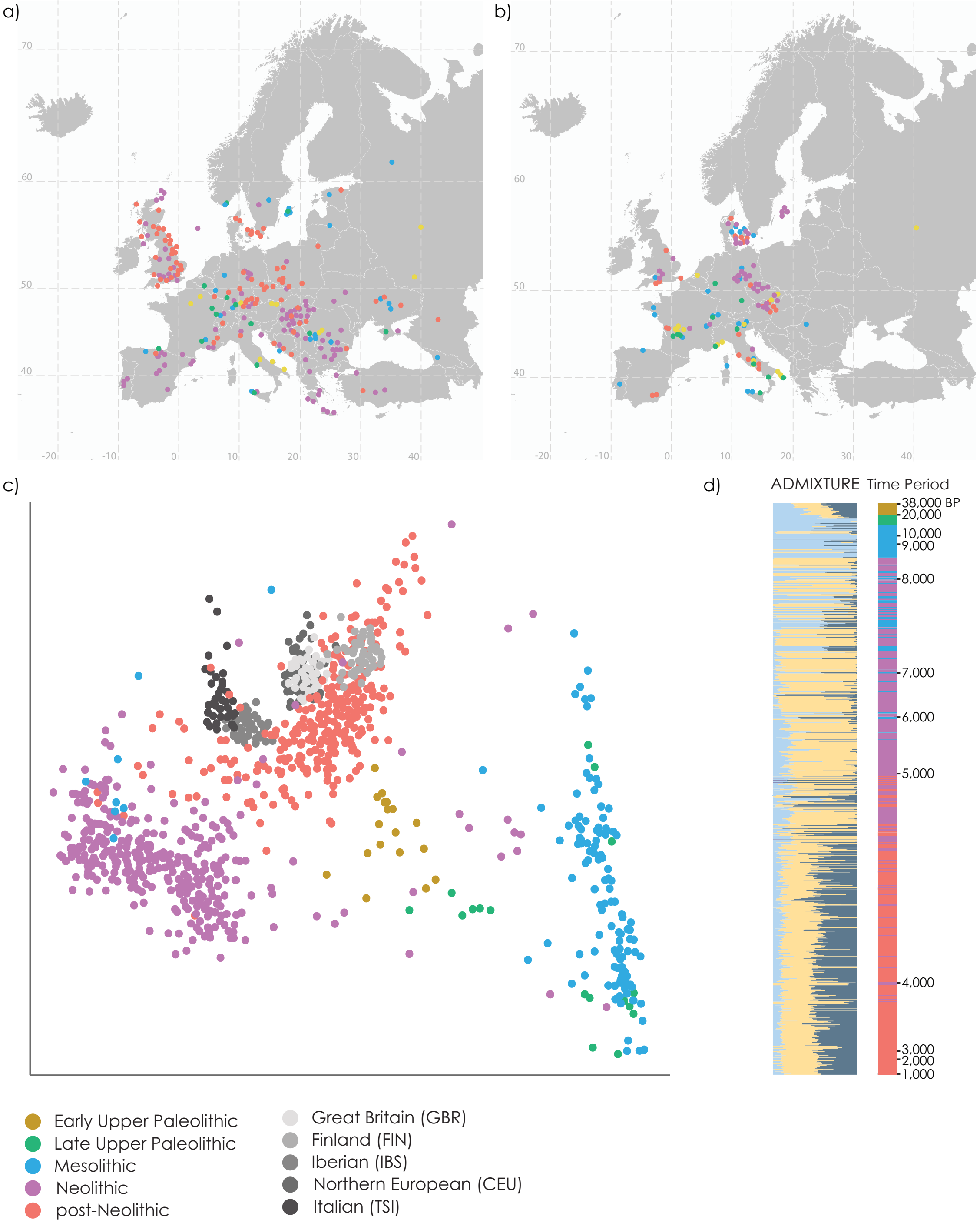


**Supplementary Figure 1:** Locations of samples, colored by time period. **a**) ancient DNA samples; **b**) skeletal samples. **c**) Two-dimensional multidimensional scaling plot of pairwise distances between ancient individuals (excluding samples with more than 99% missing data) and present-day European individuals from the 1000 Genomes Project. **d**) Unsupervised ADMIXTURE analysis of ancient individuals with k=3 (lowest cross-validation error between k=2 to 9). Rightmost panel shows population assignments.

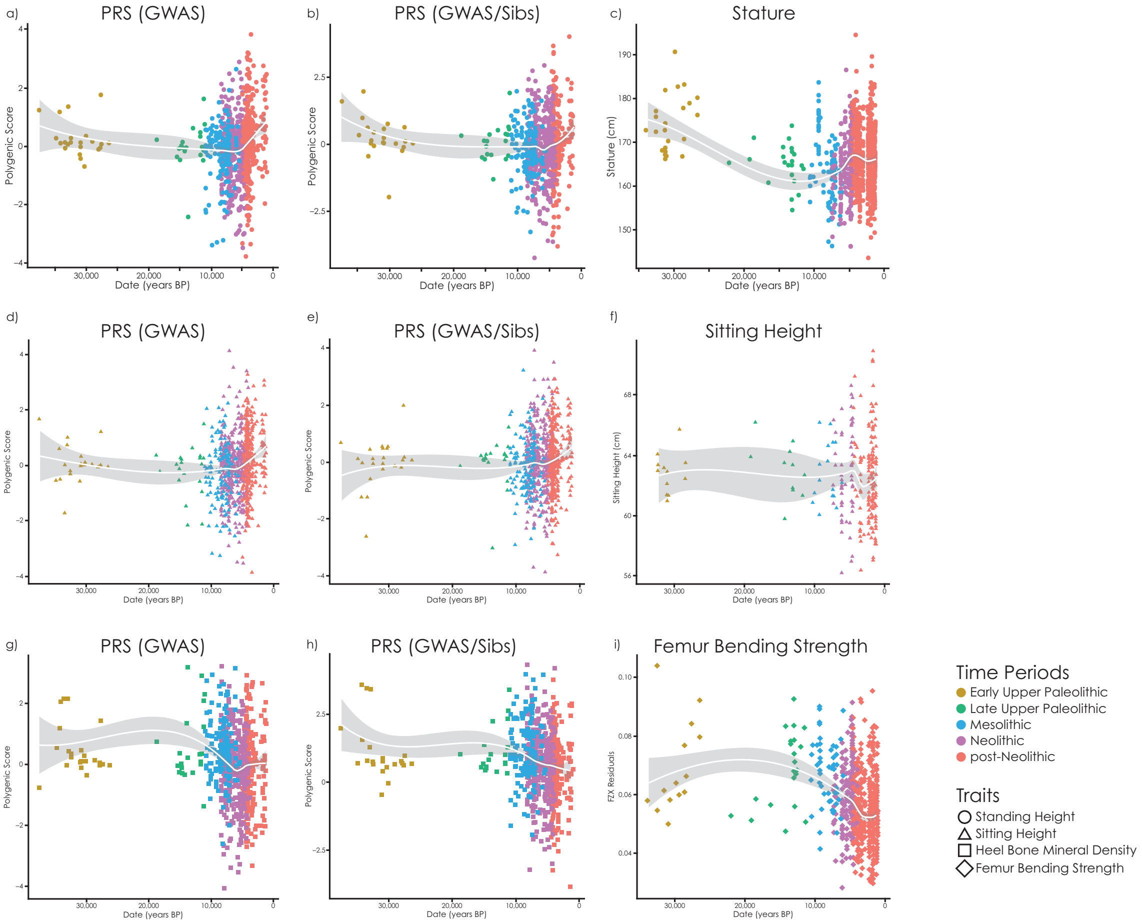


**Supplementary Figure 2:** Changes in PRS and skeletal phenotypes through time, fitting a LOESS smoothed curve to the observations without grouping. Each point is an ancient individual, lines show fitted values and grey area is the 95% confidence interval. **a**) Standing height PRS(GWAS); **b**) Standing height PRS(GWAS/Sibs); **c**) Stature (skeletal); **d**) Sitting height PRS(GWAS); **e**) Sitting height PRS(GWAS/Sibs); **f**) Sitting height (skeletal); **g**) Heel bone mineral density PRS(GWAS); **h**) Heel bone mineral density PRS(GWAS/Sibs); **i**) Femur bending strength (skeletal).

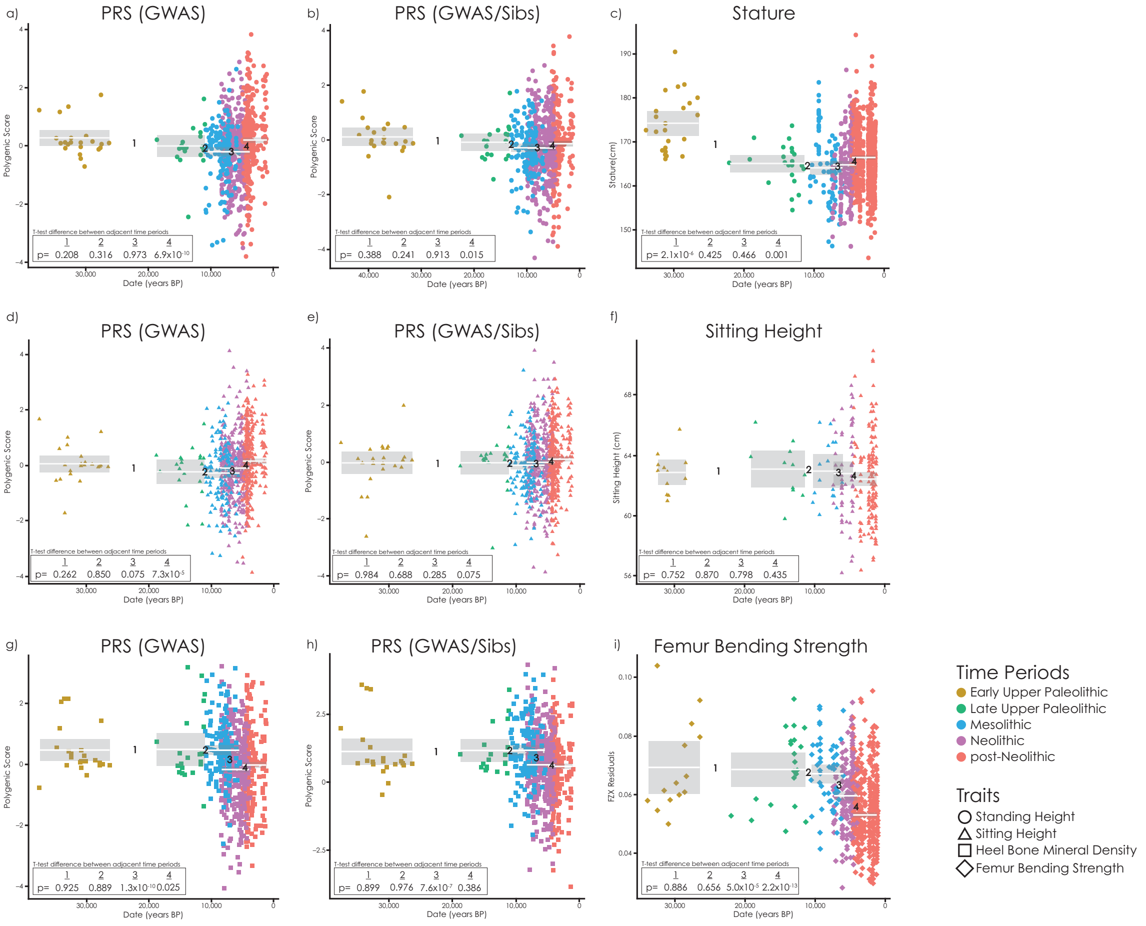


**Supplementary Figure 3:** Changes in PRS and skeletal phenotypes through time, fitting constant values in each time period. Each point is an ancient individual, lines show fitted values, grey area is the 95% confidence interval, and boxes show p-values for difference in means between adjacent groups. **a**) Standing height PRS(GWAS); **b**) Standing height PRS(GWAS/Sibs); **c**) Stature (skeletal); **d**) Sitting height PRS(GWAS); **e**) Sitting height PRS(GWAS/Sibs); **f**) Sitting height (skeletal); **g**) Heel bone mineral density PRS(GWAS); **h**) Heel bone mineral density PRS(GWAS/Sibs); **i**) Femur bending strength (skeletal).

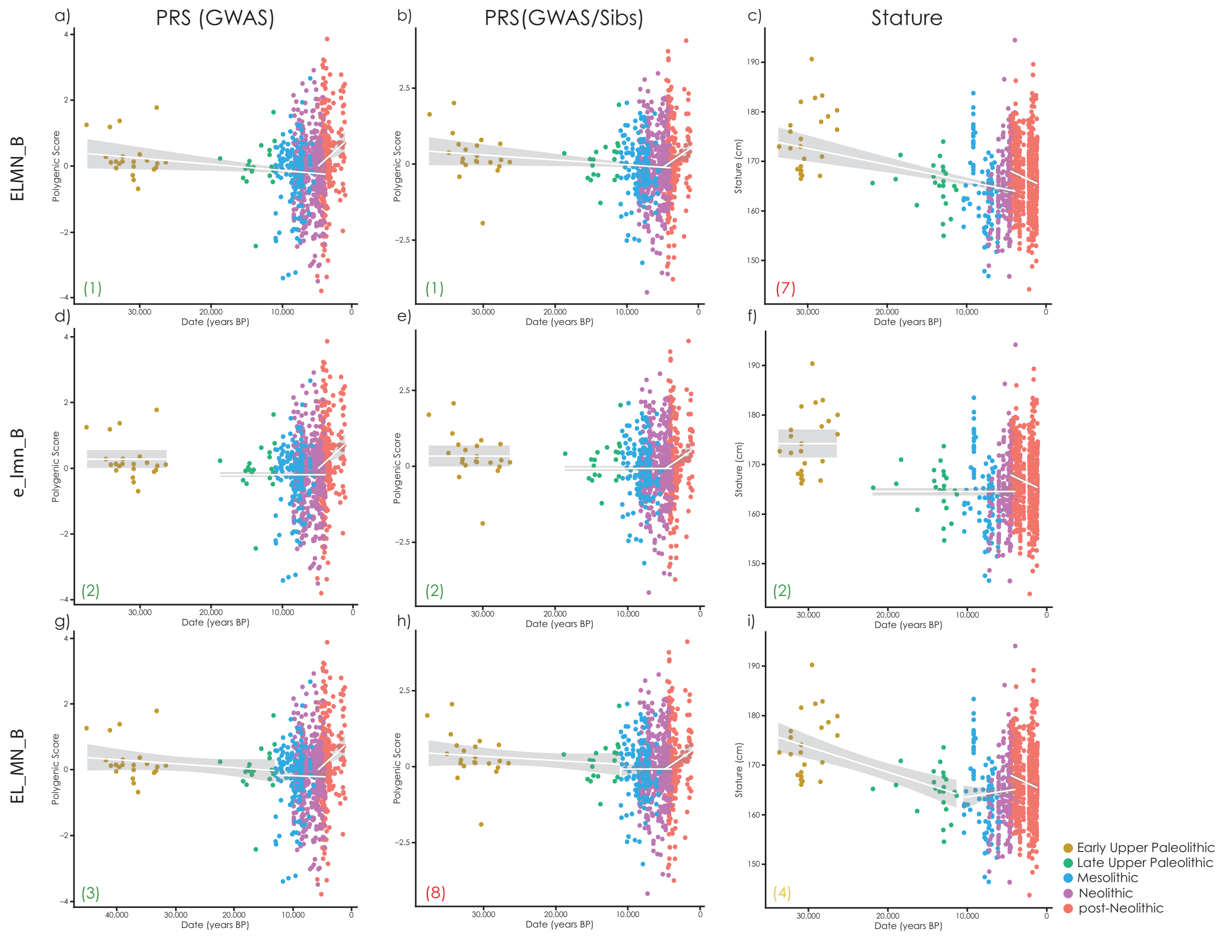


**Supplementary Figure 4:** Top three AIC models for PRS(GWAS), and the corresponding models for PRS(GWAS/Sibs) and skeletal stature. Row name indicates the model being tested, lowercase letters use fixed values for that time period, uppercase letters indicate the values were allowed to vary linearly with time. Number in the lower left corner of each plot indicates its place in the AIC ranking for PRS(GWAS), PRS(GWAS/Sibs) and Stature, green color indicates a good fit (rank 1-3/29 models), yellow a medium fit (rank 4-6/29 models), and red a poor fit (rank 7 or lower out of 29 models).

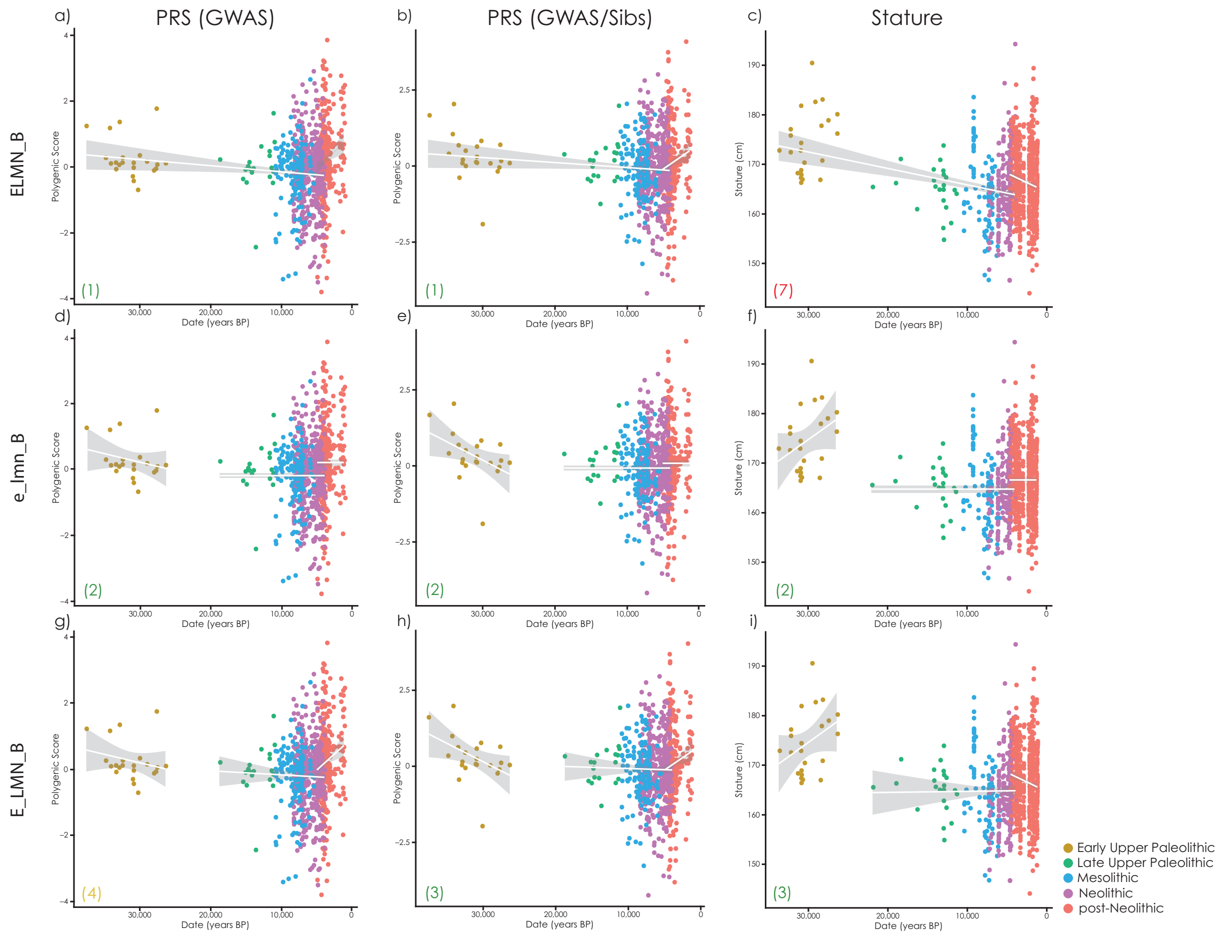


**Supplementary Figure 5:** Top three AIC models for PRS(GWAS/Sibs), and the corresponding models for PRS(GWAS) and skeletal stature. Row name indicates the model being tested, lowercase letters use fixed values for that time period, uppercase letters indicate the values were allowed to vary linearly with time. Number in the lower left corner of each plot indicates its place in the AIC ranking for PRS(GWAS), PRS(GWAS/Sibs) and Stature, green color indicates a good fit (rank 1-3/29 models), yellow a medium fit (rank 4-6/29 models), and red a poor fit (rank 7 or lower out of 29 models).

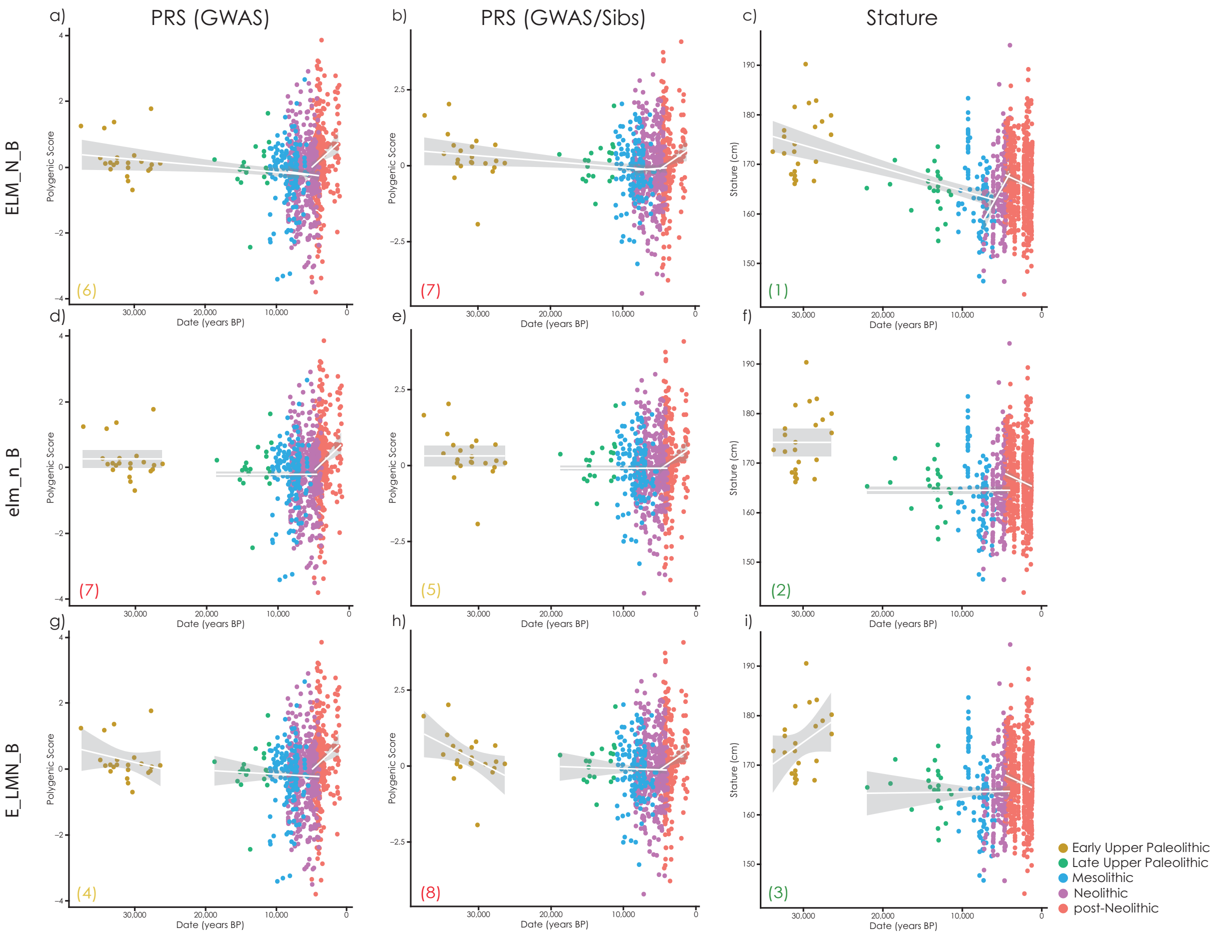


**Supplementary Figure 6:** Top three AIC models for stature, and the corresponding models for PRS(GWAS) and PRS(GWAS/Sibs). Row name indicates the model being tested, lowercase letters use fixed values for that time period, uppercase letters indicate the values were allowed to vary linearly with time. Number in the lower left corner of each plot indicates its place in the AIC ranking for PRS(GWAS), PRS(GWAS/Sibs) and Stature, green color indicates a good fit (rank 1-3/29 models), yellow a medium fit (rank 4-6/29 models), and red a poor fit (rank 7 or lower out of 29 models).

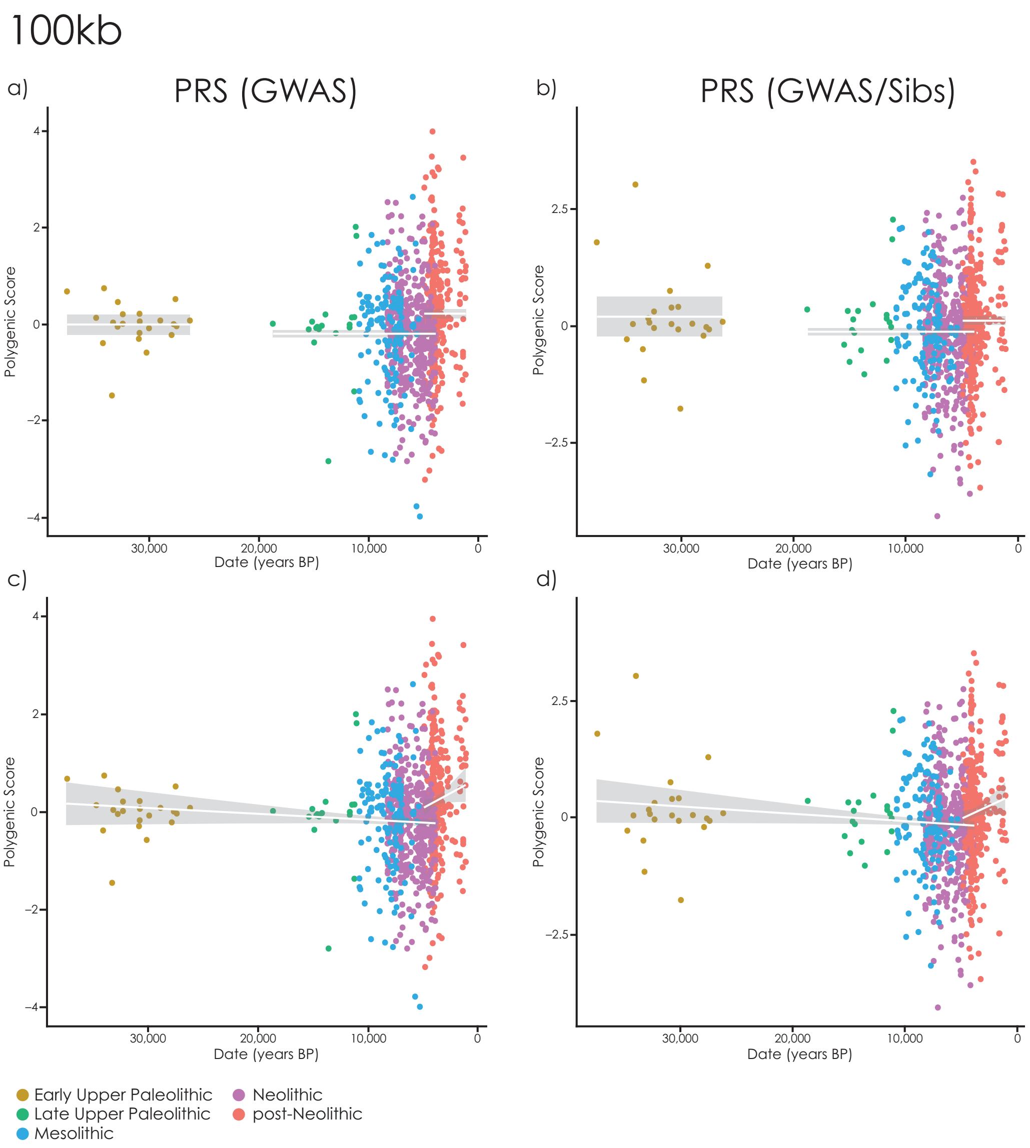


**Supplementary Figure 7:** Changes in standing height PRS though time with PRS constructed using 100kb clumping windows. Each point is an ancient individual, lines show fitted values, grey area is the 95% confidence interval. **a-b**) Constant values in the EUP, LUP-Neolithic and post-Neolithic; **c-d**) A linear trend with time between EUP-Neolithic and a different trend in the post-Neolithic.

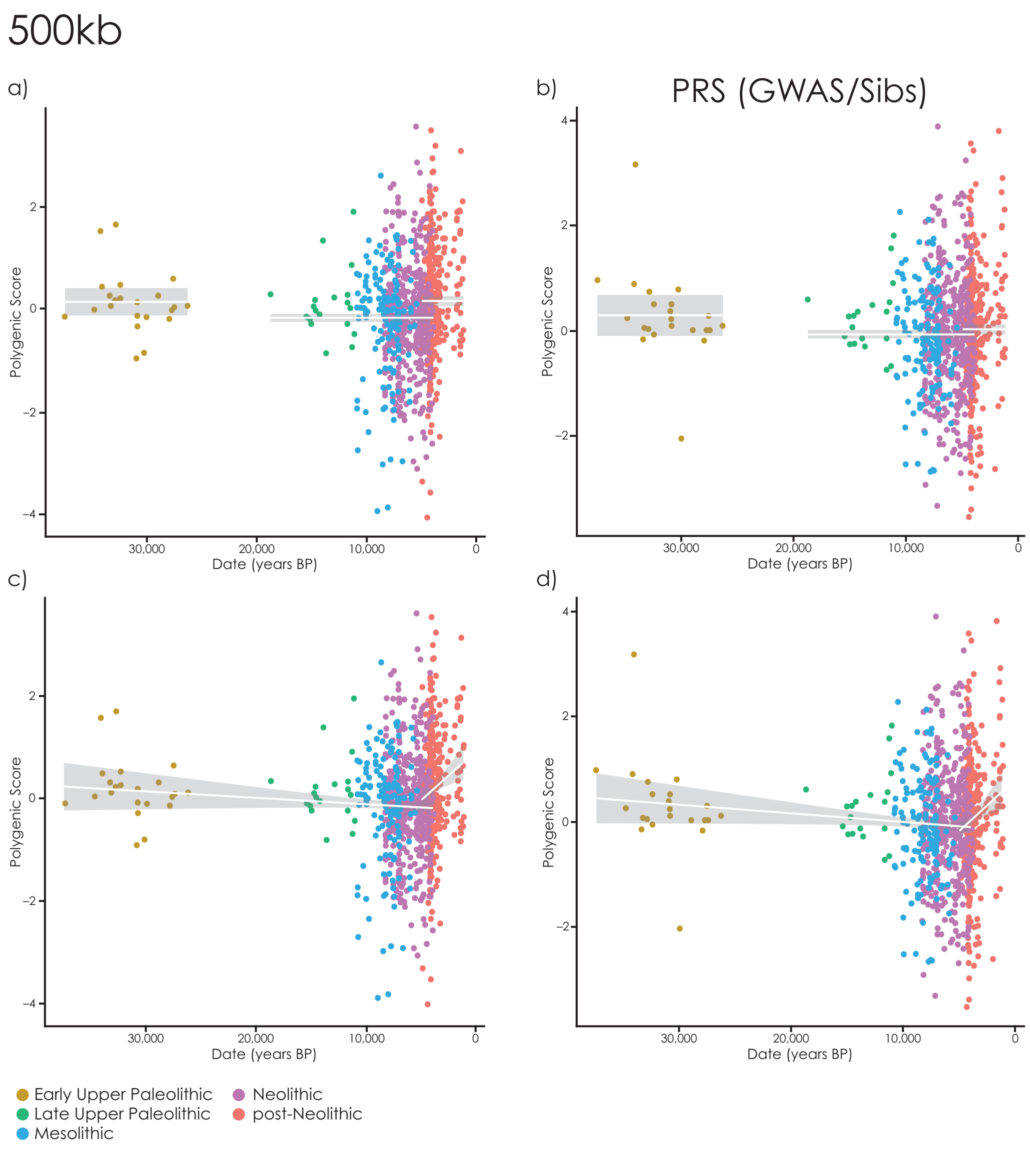


**Supplementary Figure 8:** Changes in standing height PRS though time with PRS constructed using 500kb clumping windows. Each point is an ancient individual, lines show fitted values, grey area is the 95% confidence interval. **a-b**) Constant values in the EUP, LUP-Neolithic and post-Neolithic; **c-d**) A linear trend with time between EUP-Neolithic and a different trend in the post-Neolithic.

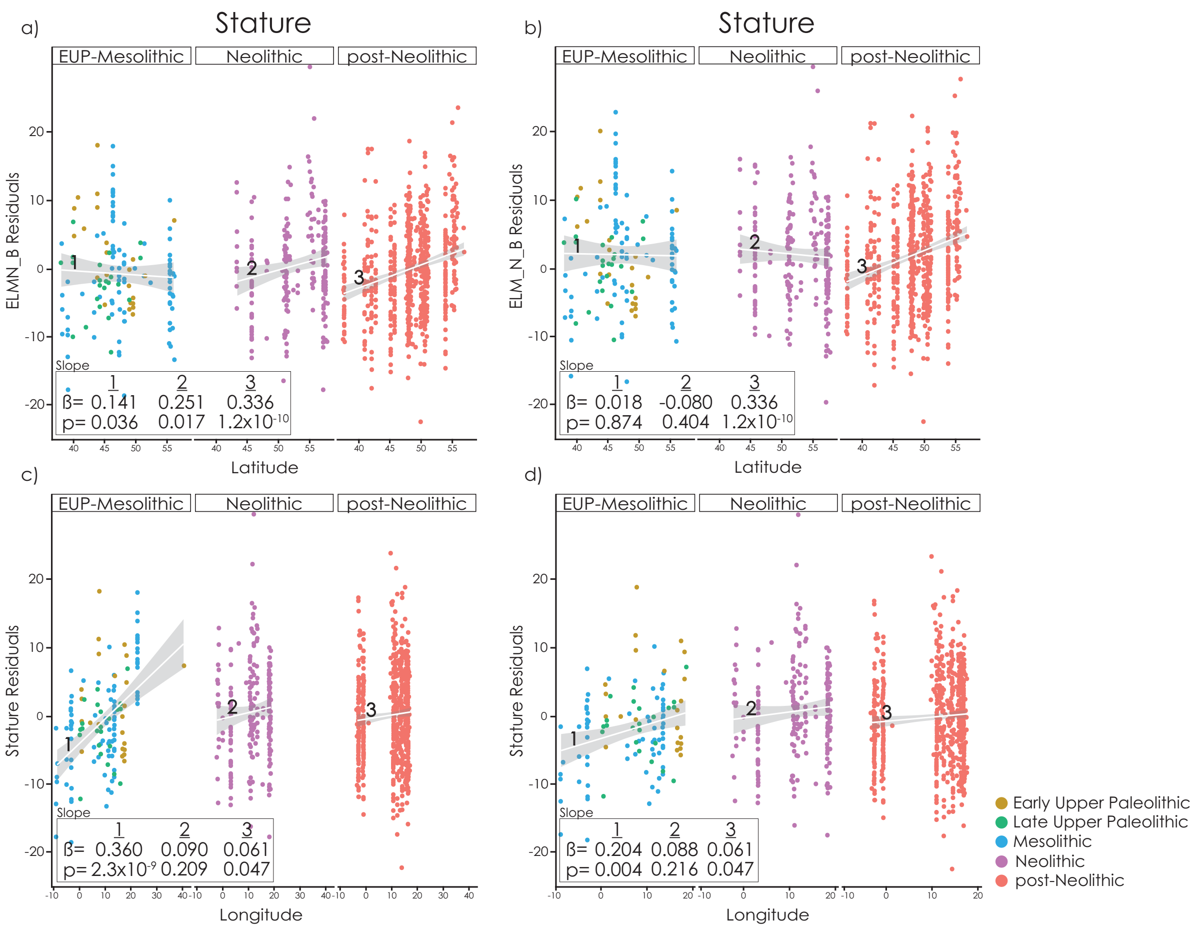


**Supplementary Figure 9:** Geographic gradients in stature under different models. Here we show EUP-Mesolithic, Neolithic and post-Neolithic periods separately, instead of EUP-Neolithic and post-Neolithic as in the main text. **a**) Latitudinal gradient using residuals of the ELMN_B model (Fig. 1f). **b**) Latitudinal gradient using residuals of the ELM_N_B model (Supplementary Fig. 5c). Note that there is no longer a gradient in the Neolithic, so the apparent geographic gradient can equally be explained by temporal change interacting with sampling. **c**) Longitudinal gradient using residuals of the ELMN_B model (Fig. 1f); the gradient is steepest in the Mesolithic and earlier. **d**) Longitudinal gradient with relatively tall Eastern Mesolithic and Paleolithic samples removed.

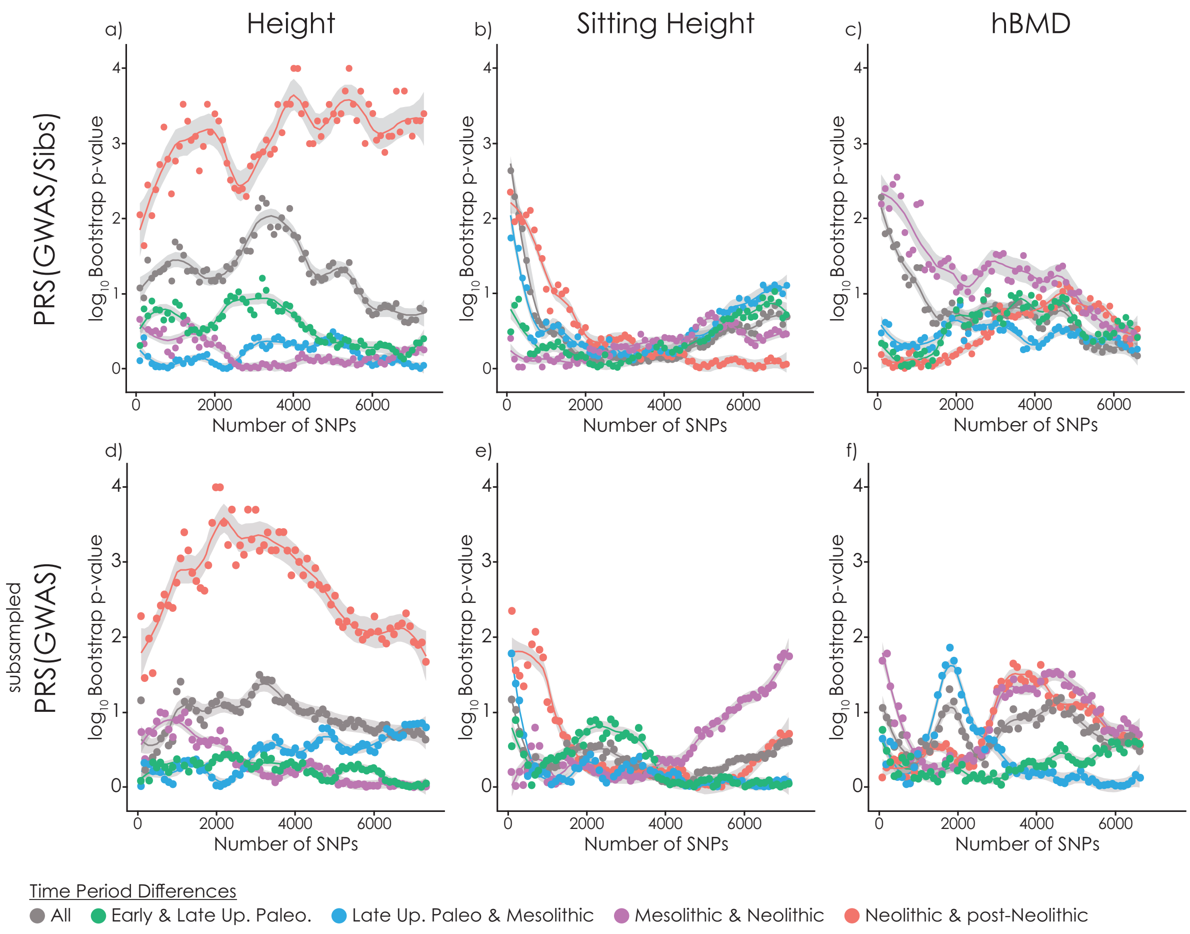


**Supplementary Figure 10**: Selection test as in Figure 6. **a-c**) Using effect sizes re-estimated within sibling pairs. **d-f**) Using GWAS results generated on a subsample of individuals so that the standard error of the effect size estimates is the same as the standard error of the within-sibling pair estimates.

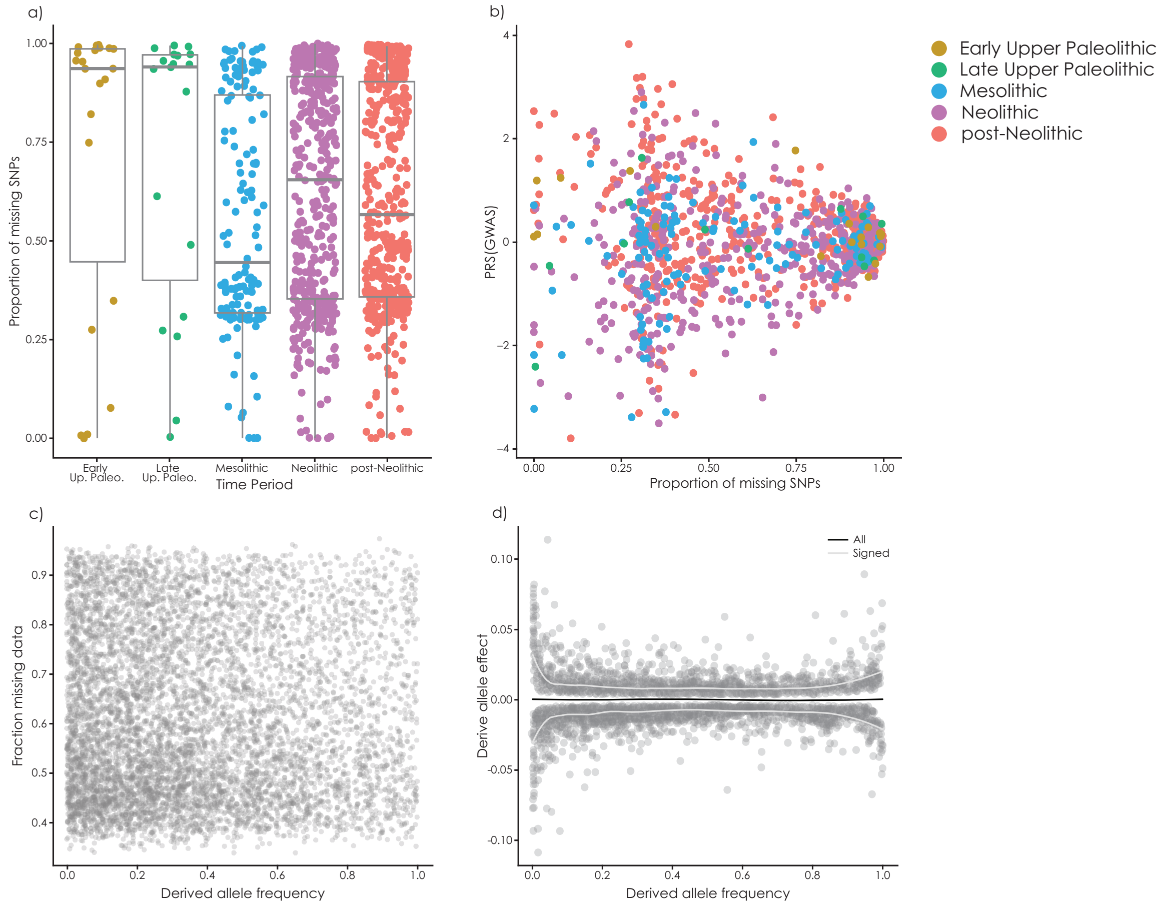


**Supplementary Figure 11:** Effect of missing data on PRS. **a**) proportion of missing data as a function of group. **b**) PRS(GWAS) as a function of missing data proportion. **c**) Missingness as a function of derived allele frequency (correlation $\rho$=0.01; not significantly different from zero, P=0.56). **d**) Derived allele effect as a function of derived allele frequency (direction of effect 3504 positive and 3529 negative; proportion positive not significantly different from 0.5, P=0.77).
